## Supplemental Material for "The architecture of EMC reveals a path for membrane protein insertion"

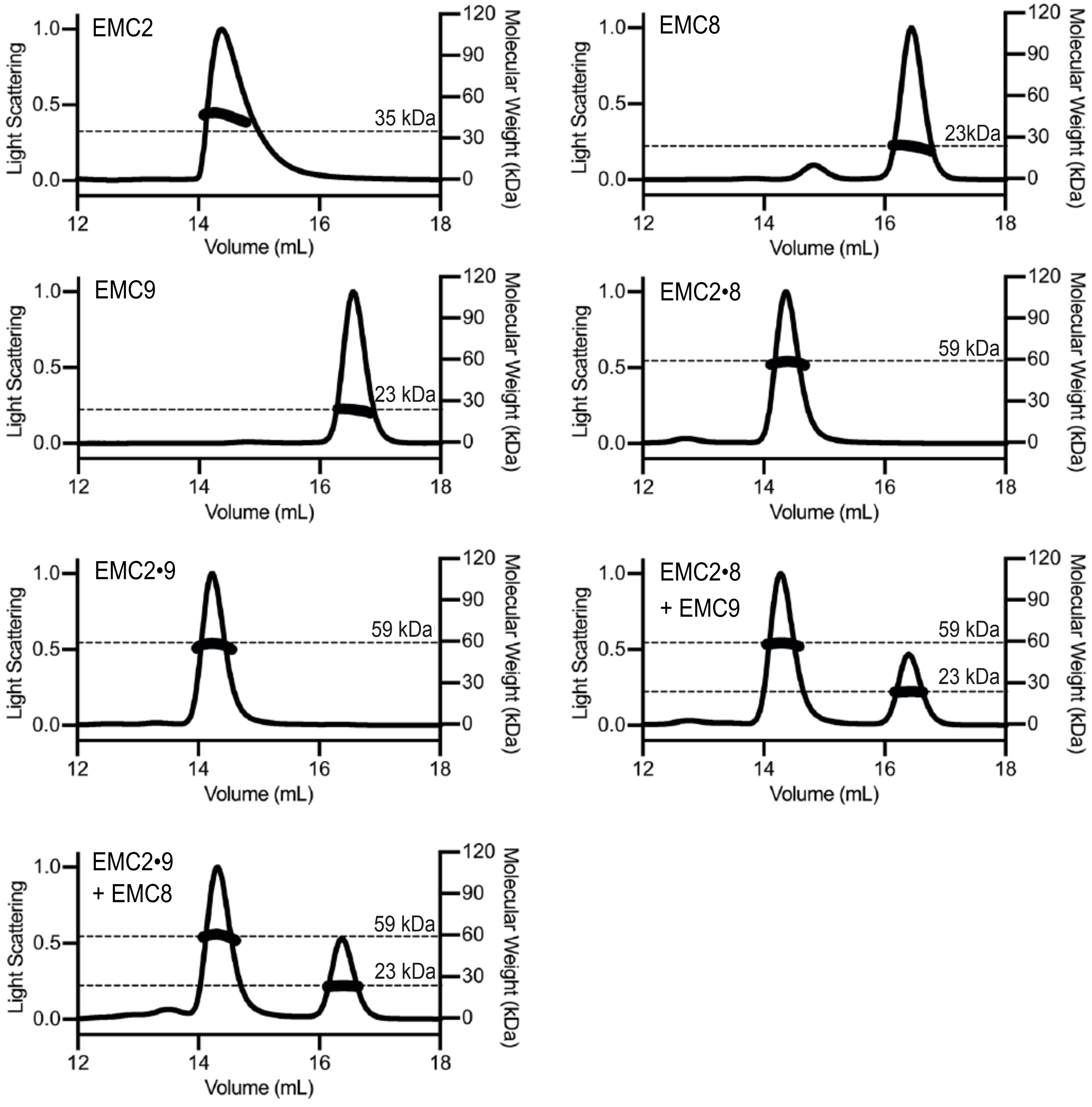


**Fig. 1 – figure supplement 1. SEC-MALS of individual and complexed cytoplasmic EMC-subunits.** Size Exclusion Multi Angle Light Scattering (SEC-MALS) analysis of recombinant EMC2, EMC8 and EMC9. The signal from the 90°-scattering detector is plotted on the left y-axis as a solid line and the calculated molecular weight is plotted on the right y-axis and depicted as points across the elution peak. A dotted line notated with text indicates the theoretical molecular weight for each protein. The calculated molecular weights are consistent with those predicted for monomeric subunits and complexed subunits. Neither addition of EMC8 to a preformed EMC2•9 complex nor addition of EMC9 to a pre-formed EMC2•8 complex resulted in the formation of a ternary complex.

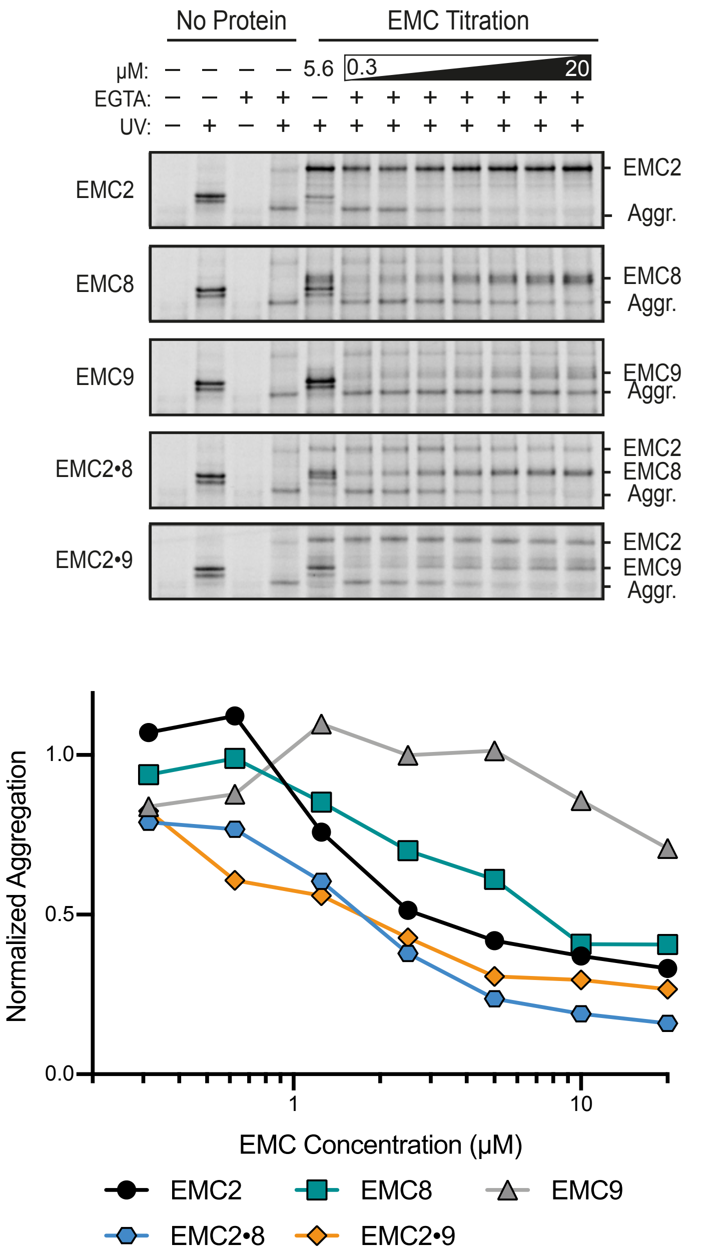


**Fig. 1 – figure supplement 2. Activity of EMC cytosolic subunits in preventing TMD aggregation.** $^{35}$S-methionine labeled SQS containing the benzoyl-phenylalanine (Bpa) photo-crosslinker within the TMD was prepared in complex with CaM as characterized in Fig. 1C. The complex was mixed with various concentrations of different proteins (indicated on the left), treated with EGTA to release the substrate from CaM, and substrate interactions monitored by UV-mediated crosslink formation. Release from CaM in the absence of any additional protein results in UV-dependent self crosslinks indicating of aggregation (Aggr.). This aggregate product is progressively reduced to a comparable extent by the inclusion of EMC2, EMC2•8, or EMC2•9. By contrast, EMC9 alone is not able to reduce aggregation, while EMC8 alone has a very modest effect on aggregation. Note that the crosslinking observed to EMC8 or EMC9 when they are added in isolation is probably due to their co-aggregation with substrate. The plot below the gels quantifies the relative intensity of the aggregate band normalized to the sample without added protein.

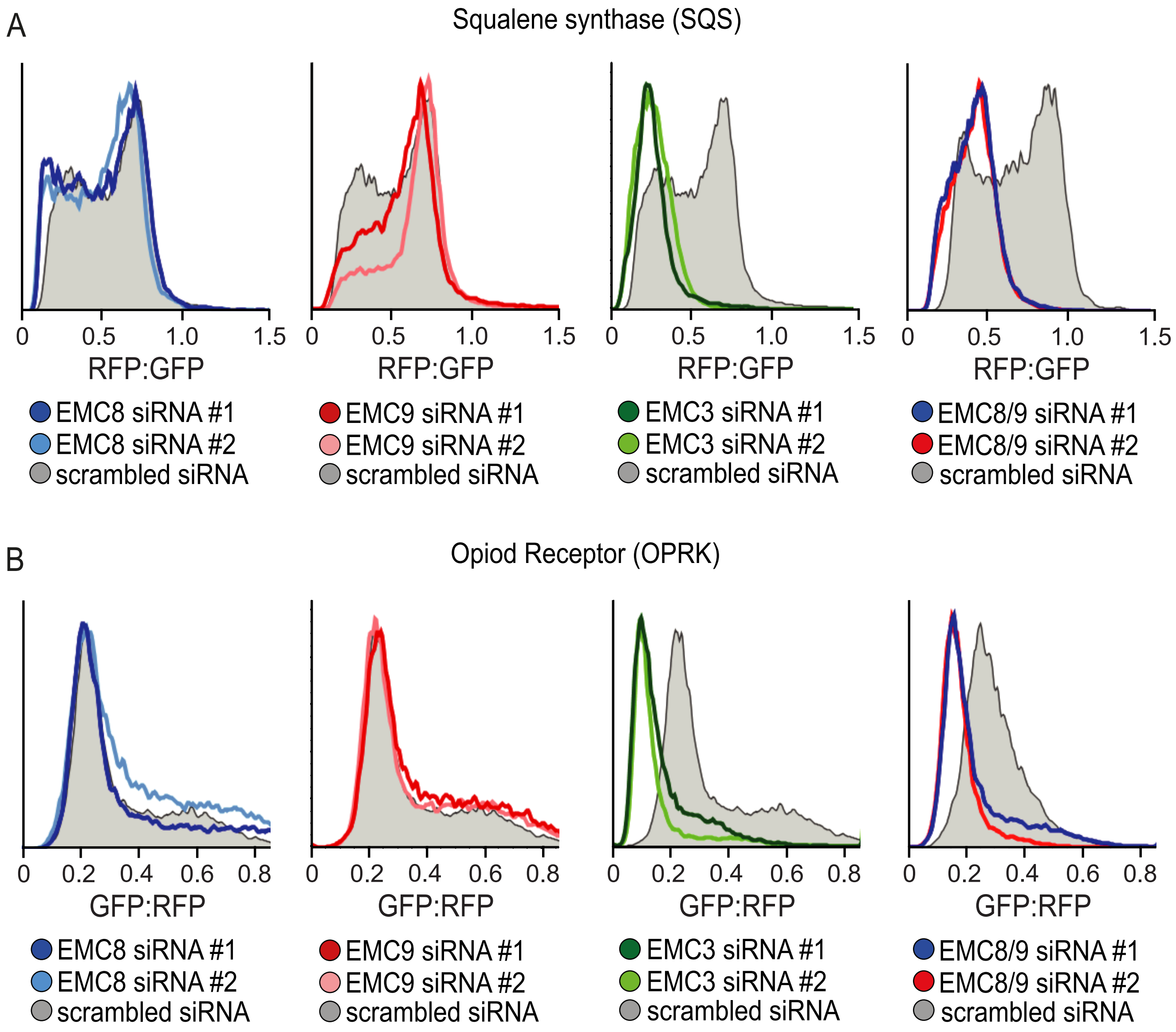


**Fig. 1 – figure supplement 3. EMC8 and EMC9 necessary for membrane protein biogenesis but are functionally redundant.** (A) Flow cytometry analysis of a tail-anchored RFP-squalene synthase (RFP-SQS) substrate, relative to an internal GFP expression control. Reduced biogenesis of the TMD substrate results in a shift of the histogram ratio to the left. siRNA mediated knockdown of either EMC8 or EMC9 had minimal impact on RFP-squalene synthase biogenesis compared to knockdown of the integral transmembrane component EMC3. Combined knockdown of EMC8 and EMC9 resulted in a significant defect in RFP-SQS biogenesis. (B) The same experiment as in panel A but using the mu opioid receptor (GFP-OPRK), a multi-pass membrane protein, as the substrate. Here, OPRK is GFP-tagged and RFP serves as the internal expression control.

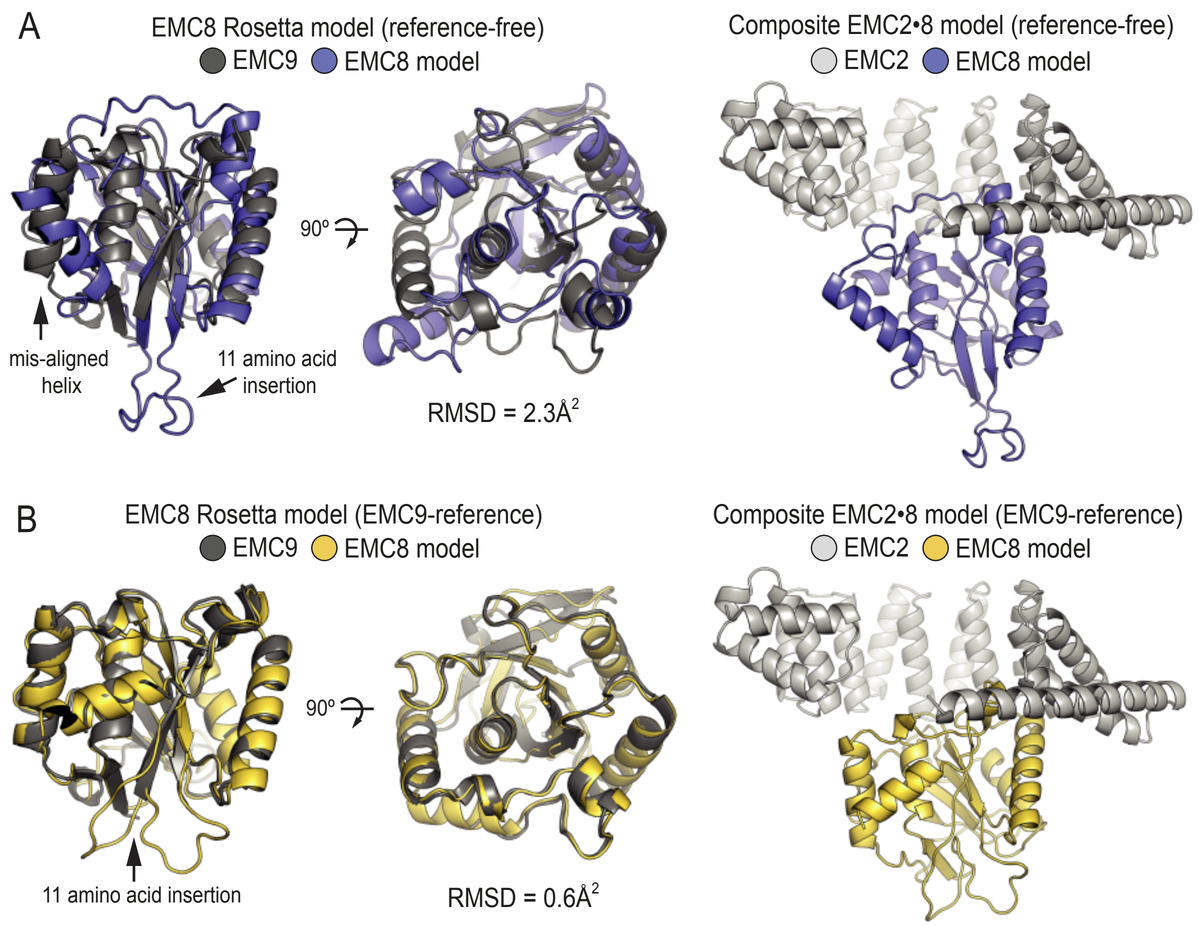


**Fig. 2 – figure supplement 1. Protein structure prediction confirms structural homology between EMC9 and EMC8.** (A) Reference free ab initio modelling of EMC8 (blue) using Robetta results in a model with an overall RMSD of 2.3 $Å^2$ when aligned to the EMC9 crystal structure (grey). The majority of this disparity is the result of a single misaligned helix and the location of an 11 residue insertion in a loop in EMC8. Docking of the predicted structure into the EMC2•9 structure in the place of EMC9 results in a good fit with no steric violations. (B) Ab initio modelling of EMC8 (yellow) using Robetta with the EMC9 crystal structure (grey) as a template results in an improved model with an overall RMSD of 0.6 $Å^2$. As with the reference-free model, the template-based model of EMC8 yields the same core fold and fits well when substituted into the EMC2•9 crystal structure.

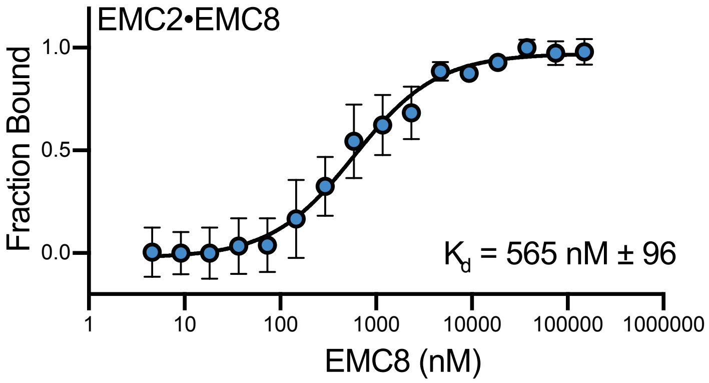


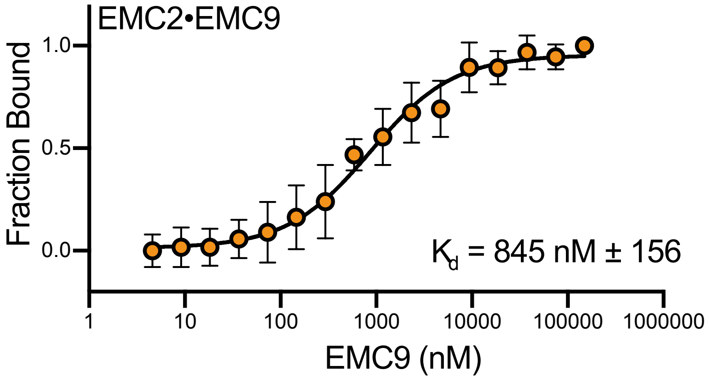


**Fig. 2 – figure supplement 2. EMC8 and EMC9 have similar affinities for EMC2.** Microscale thermophoresis was used to determine affinities of the heterodimer. An endogenous solvent accessible cysteine in EMC2 was labeled with maleimide-OG488 and held at a fixed concentration of 50 nM. Either EMC8 or EMC9 were titrated from 4 nM - 150 µM. Affinities calculated for EMC2•8 (blue) and EMC2•9 (orange) were ~550nM and ~850nM respectively. Thermophoresis data were fit using NanoTemper's in software quadratic model, and equivalent results were attained using nonlinear fit model in Prism GraphPad. Data consist of two biological replicates with two technical replicates each. Data points represent the mean and error bars represent standard deviation.

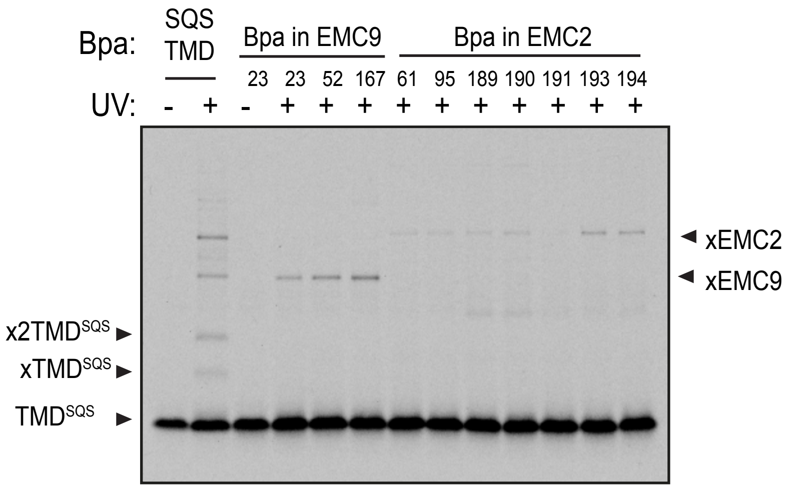


**Fig. 3 – figure supplement 1. Crosslinking analysis of the EMC2•9 cytosolic vestibule.**
$^{35}$S-methionine-labeled TMD of SQS was mixed with recombinant purified EMC2•9 complex as in Fig. 1 and analyzed directly or after UV irradiation as indicated. The photocrosslinking amino acid benzoyl-phenylalanine (Bpa) was incorporated into either the SQS substrate, EMC9, or EMC2 as indicated.

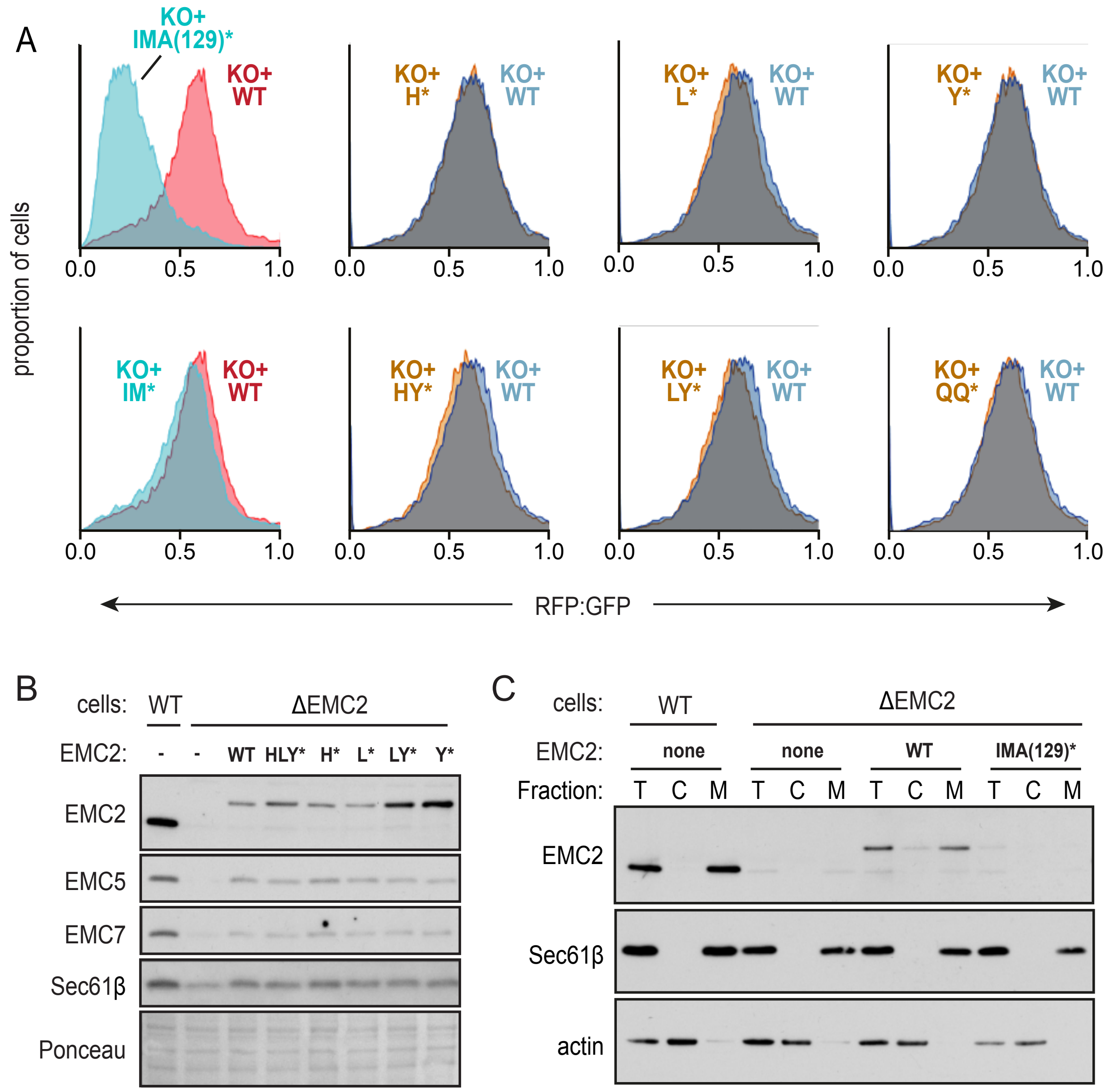


**Fig. 3 – figure supplement 2. Functional analysis of the EMC2•9 cytosolic vestibule.** (A) Expression of the dual color SQS reporter in EMC2 knockout cells complemented with either WT EMC2 or the indicated EMC2 mutants. The data are represented as histograms of the RFP to GFP ratio. The two left graphs are from a different experiment from the remainder of graphs (which were analyzed together) and are colored differently to indicate this. The knockout-like phenotype of the IMA(129)* mutation is apparently due to the A129K mutation since the IM* mutation shows almost complete rescue. The mutated amino acids are: I61K, M95K, A129K, H189E, L190E, Y191K, Q193E, and Q194K. (B) Immunoblotting of whole cell extracts from WT or ΔEMC2 cells transfected with the indicated EMC2 constructs. A section of the Ponceau-stained blot shows equal loading. (C) WT and ΔEMC2 cells transfected with the indicated EMC2 constructs were fractionated into cytosol (C) and membrane (M) fractions and analyzed by immunoblotting for EMC2 relative to an equivalent amount of total (T) cell lysate. The membrane marker Sec61β and cytosolic marker actin were included as controls. Note that relative to WT EMC2 expressed in ΔEMC2 cells or endogenous EMC2 in WT cells, the IMA(129)* mutant is very poorly expressed. This is presumably due to its degradation secondary to inefficient assembly with other EMC subunits, explaining its strong phenotype in the flow cytometry assay.

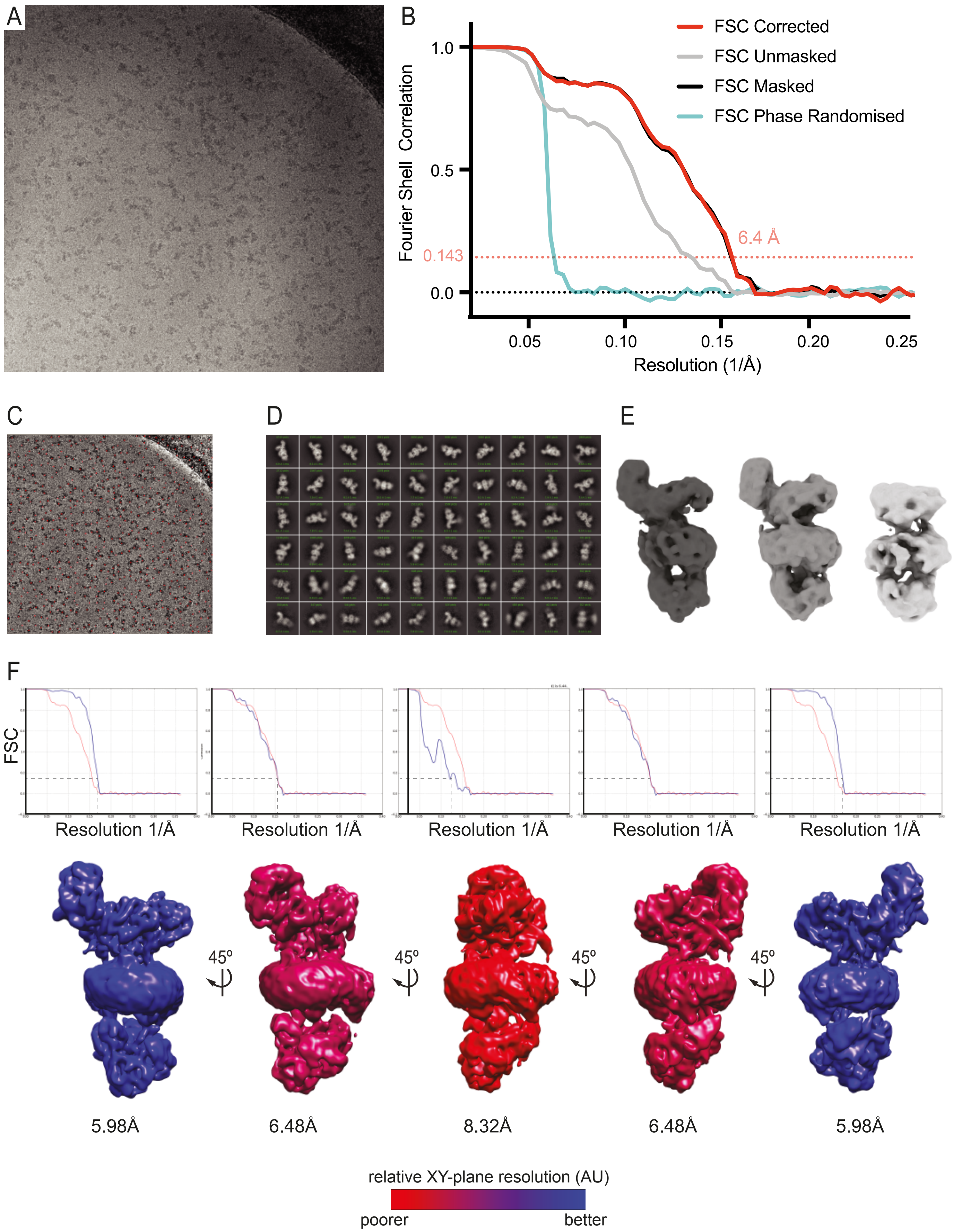


**Fig. 4 – figure supplement 1. Cryo-electron microscopy data processing.** (A) Micrograph showing EMC particles imaged at -0.5 µm defocus with volta phase plate at a nominal pixel size of 1.38 Å using a K2 summit detector. (B) Fourier Shell Correlation (FSC) curves for phase randomised, unmasked, masked and corrected 3D reconstructions. Representative images of template-based autopicking (C), 2D-classification (D) and 3D classification (E) stages during the processing of dataset 1. (F) 3D FSC curves demonstrating resolution anisotropy during reconstruction of a map of the human EMC. 5 views of the same map are shown in rotational increments of 45º. Each view is accompanied by an FSC plot comparing the FSC in the respective viewing orientation (blue) to the global FSC (red). Resolution values given below the maps are calculated from the FSC plots above the maps using the 0.143 criterion. Maps are coloured from blue (high) to red (low) to indicate reconstruction resolution from different angles. Maps generated using 3DFSC implemented in CryoSPARC (Tan et al., 2017).

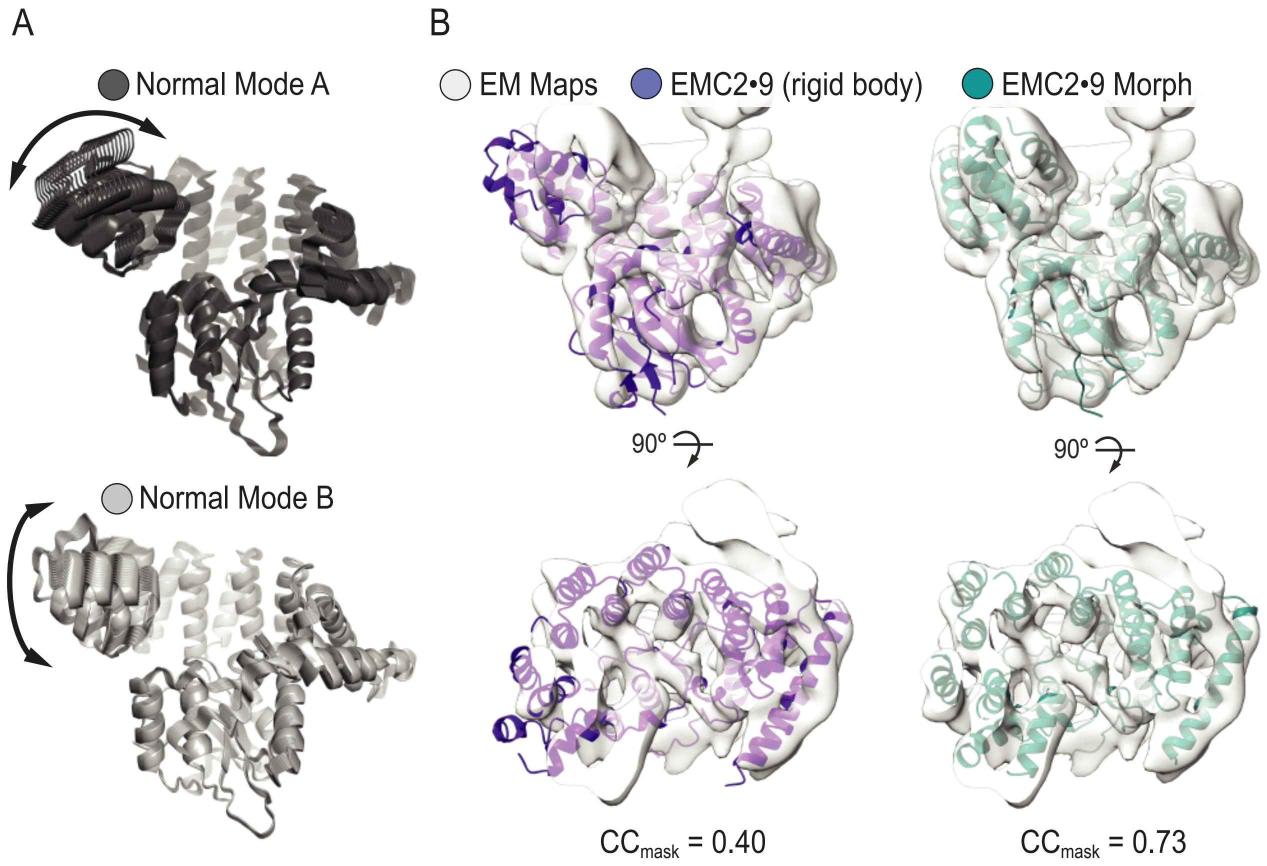


**Fig. 4 – figure supplement 2. Normal mode analysis and flexible fitting of the EMC2•9 crystal structure into the full EMC cryo-EM map.** (A) Normal mode analysis using the el Némo server (Suhre and Sanejouand, 2004) reveals conformational flexibility in the N-terminal region of EMC2•9. All calculated states are shown for modes A and B in dark and light grey, respectively. Arrows indicate direction of motion. (B) Rigid body docking of the EMC2•9 crystal structure (purple) into the experimental cryo-EM density (grey) is in overall good agreement with the exception of EMC2's N-terminal region. The fit is improved (teal) following flexible fitting conducted with Flex-EM, and real space refinement in Coot and PHENIX. The map to model CCmask calculated in PHENIX real space refine increases from 0.40 for the rigid body fit to 0.73 for the flexibly fitted structure. During flexible fitting the largest movements occurred in the N-terminal helical bundle of EMC2, consistent with the normal mode analysis.

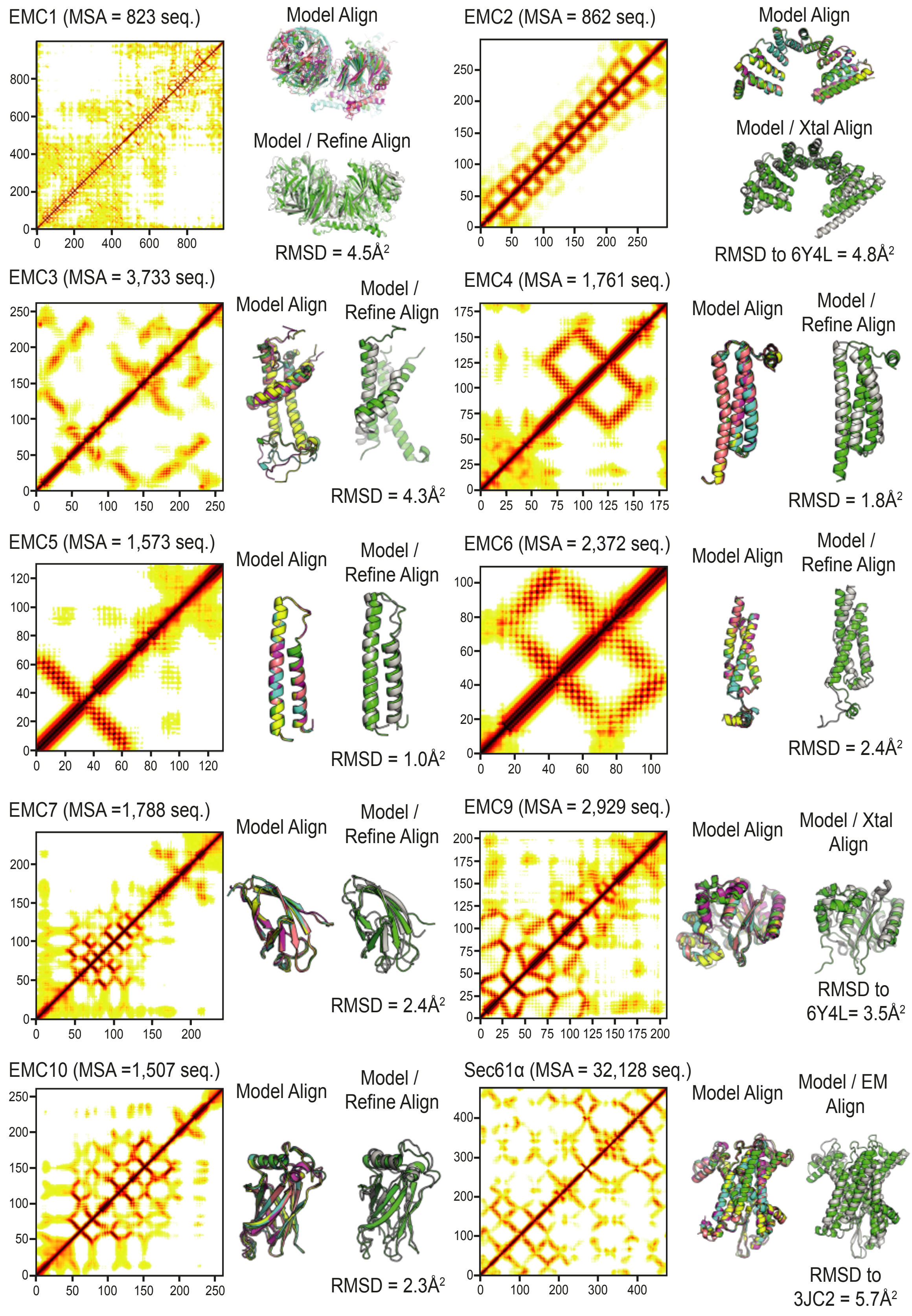


**Fig. 5 – figure supplement 1. Ab initio prediction of EMC subunit structure and flexible fitting using real-space refinement.** Structures were predicted for 9 EMC subunits using the trRosetta server. The best ranked solution for each subunit was then docked into the cryo-EM density and subject to real-space refinement in PHENIX. Number of alignments in the multiple sequence alignment (MSA) is indicated along with the resulting distance contact heat map. The top 5 ranked models were aligned for the core folds used in EM docking to show modelling confidence. A comparison of the model before and after real space refinement illustrates only subtle changes (green vs grey, RMDS is indicated). Soluble and membrane proteins (EMC2, EMC9, and Sec61α) were modelled and compared to their known structures.

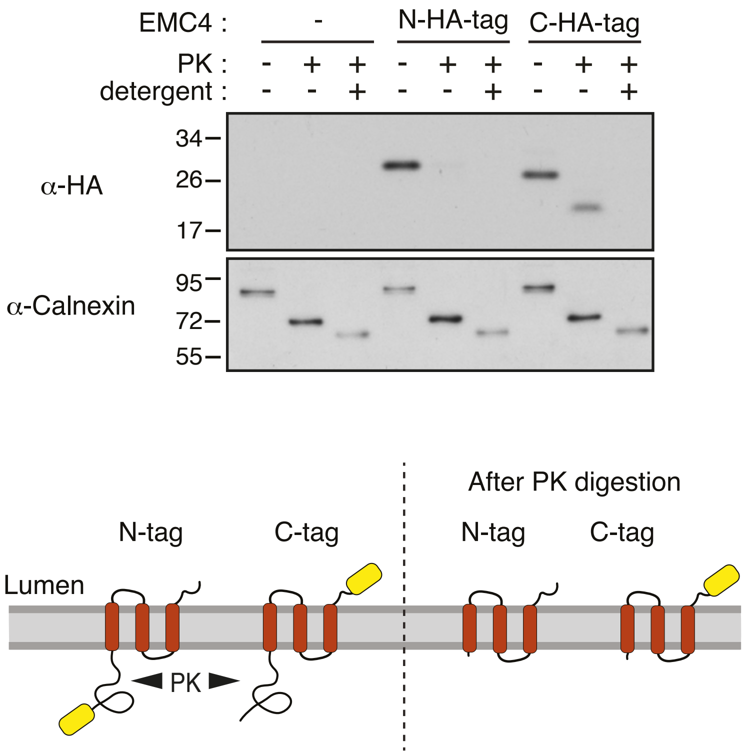


**Fig. 5 – figure supplement 2. Protease-protection analysis of EMC4 topology.** Microsomes isolated from HEK293T cells transfected with an empty vector, EMC4 with an N-terminal 3xHA tag (N-HA-tag), or EMC4 with a C-terminal 3xHA tag (C-HA-tag) were analyzed by a protease-protection assay. Equal aliquots were left untreated, treated with proteinase K (PK), or treated with PK in the presence of detergent (Triton X-100) and analyzed by immunoblotting for the HA tag and the N terminus of calnexin. The N-terminal 3xHA tag was accessible to PK in the absence of detergent, suggesting the N-terminus is exposed to the cytosol. The C-terminal 3xHA tag was protected from PK digestion in the absence, but not presence, of detergent, suggesting that the C-terminus is in the ER lumen. This supports a three TMD model in which the loop between TMD2 and TMD3 is too short to be accessible to PK. A schematic of the results and proposed topology of EMC4 is also shown. In principle, the protease-digestion results are also consistent with a single-spanning topology with the N-terminus facing the cytosol. However, this is unlikely given that all topology prediction algorithms predict at least two TMDs and trRosetta predicts a three helix bundle. Note that the core region in the N-terminal lumenal domain of calnexin is resistant to complete PK digestion.

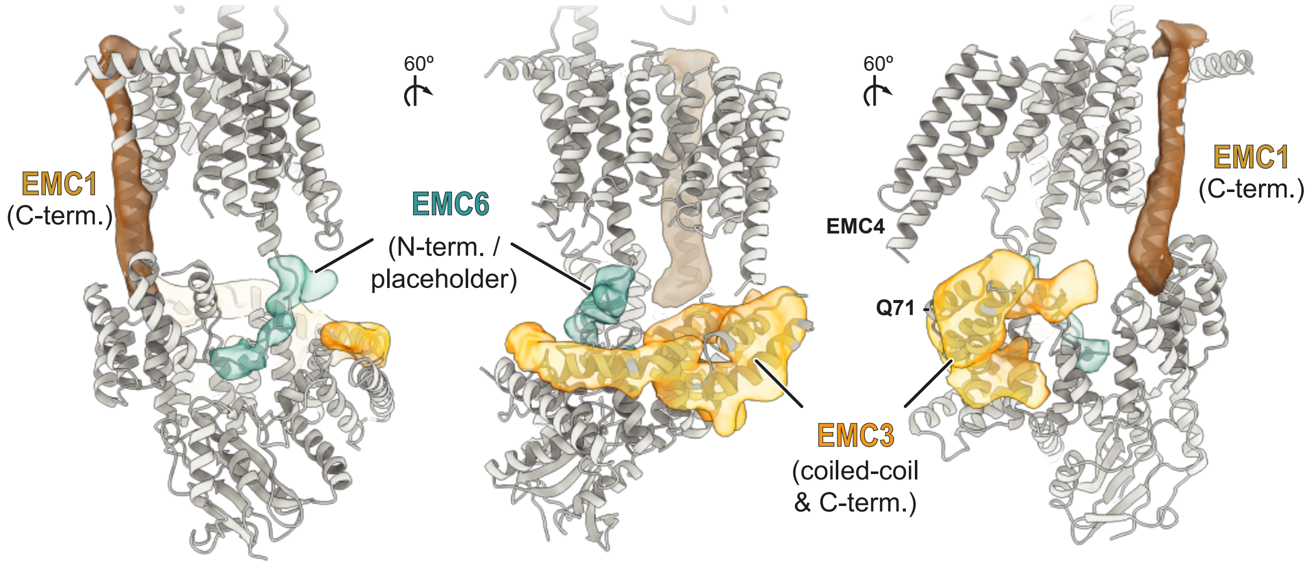


**Fig. 5 – figure supplement 3. Provisional assignment of non-EMC2•9 cytosolic density.** Semi-automated segmentation in UCSF chimera revealed regions of density in the cytosolic domain of the cryo-EM map of the EMC which are not accounted for by the EMC2•9 heterodimer (translucent densities). Based on the position and connectivity to the respective TMDs in the membrane portions of this density were provisionally assigned to EMC1 (brown) and the N-terminus of EMC6 (teal) which is proposed to form a 'placeholder' in the cytosolic substrate binding pocket. A predicted three-helix bundle formed from a coiled-coil between TMD1 and TMD2 of EMC3 interacting with a helix in EMC3's C-terminal tail was fit into a similarly shaped and sized density. In this fitted model, Q71 of EMC3 is within 15 Å of the position assigned to EMC4. AbK incorporated at Q71 shows a UV-dependent crosslink to EMC4, consistent with our assignments. The remaining density is continuous with the C-terminal helix in this three-helix bundle, and was therefore provisionally assigned to the rest of EMC3's C-terminal tail.

| Table S1: X-ray data collection and refinement statistics | | |
|---|---|---|
| | EMC2•9<br>(S-SAD phasing) | EMC2•9<br>(Molecular replacement) |
| **Data collection** | | |
| X-ray source | Diamond I23 | Diamond I03 |
| X-ray wavelength (Å) | 2.7552 | 0.9763 |
| Space group | $P2_12_12_1$ | $P2_12_12_1$ |
| Unit cell parameters | | |
| a, b, c (Å) | 53.2 82.8 124.0 | 52.6 84.9 122.6 |
| α, β, γ (º) | 90.0 90.0 90.0 | 90.0 90.0 90.0 |
| Resolution range (Å) | 49.6 - 2.65 (2.78 - 2.65) | 49.7 - 2.20 (2.27 - 2.2) |
| No. of reflections | | |
| Total | 668148 (73113) | 1486886 (128644) |
| Unique | 16468 (2109) | 28668 (2433) |
| Completeness (%) | 99.7 (98.9) | 100.0 (100.0) |
| Multiplicity | 40.6 (34.7) | 51.9 (52.9) |
| I/σ(I) | 36.8 (1.8) | 17.6 (1.9) |
| $R_{meas}$ (%) | 6.4 (214.6) | 17.2 (465.7) |
| $R_{merge}$ (%) | 6.3 (211.5) | 17.0 (461.3) |
| $R_{pim}$ (%) | 1.0 (35.3) | 2.4 (63.7) |
| $CC_{1/2}$ (%) | 100.0 (81.7) | 100.0 (92.6) |
| **Refinement** | | |
| $R_{work}$ / $R_{free}$ (%) | - | 20.3 / 25.0 |
| RMS deviations | | |
| Bond length (Å) | - | 0.007 |
| Bond angle (º) | - | 0.848 |
| No. of atoms | | |
| Protein | - | 3454 |
| Ligands | - | 72 |
| Water | - | 80 |
| Average B-factors (Å$^2$) | | |
| Total | - | 70.6 |
| Protein | - | 71.7 |
| Ligands and waters | - | 65.4 |
| Ramachandran (%) | | |
| Favored | - | 96.4 |
| Outliers | - | 0.0 |
| PDB Code: | - | 6Y4L |

(*) Values in brackets are for the highest resolution bin

Table S2: EM collection and processing

| **Collection Parameters** | **Dataset 1** | **Dataset 2** | **Dataset 3** | **Dataset 4** | **Dataset 5** |
|---|---|---|---|---|---|
| Microscope | Titan Krios (m06 eBIC) | Titan Krios (m06 eBIC) | Titan Krios (m06 eBIC) | Titan Krios (MRC-LMB) | Titan Krios (MRC-LMB) |
| Pixel Size | 1.380 | 1.380 | 1.380 | 1.179 | 1.390 |
| Voltage | 300 | 300 | 300 | 300 | 300 |
| Spherical Aberation | 2.7 | 2.7 | 2.7 | 2.7 | 2.7 |
| Total exposure ($e^-$/Å$^2$) | 39.60 | 42.50 | 37.77 | 39.36 | 44.36 |
| Exposure Length (s) | 5.0 | 11.02 | 14 | 11 | 11 |
| Frames | 25 | 44 | 40 | 44 | 40 |
| Defocus Range (μm) | -0.5 to -1.5 | -0.5 to -1.5 | -0.5 to -1.5 | -0.5 to -1.5 | -0.5 to -1.5 |
| Micrographs | 2776 | 2484 | 1206 | 4228 | 932 |
| Microscope tilt (degrees) | 0 | 0 | 30 | 20 | 20 |
| Volta Phase Plate | ✓ | ✓ | ✓ | ✓ | ✓ |

| **Pre-merge processing** | **Dataset 1** | **Dataset 2** | **Dataset 3** | **Dataset 4** | **Dataset 5** |
|---|---|---|---|---|---|
| Motion Correction & CTF estimation | 2776 micrographs | 2484 micrographs | 1206 micrographs | 4228 micrographs | 932 micrographs |
| Blob-based autopicking | 1,298,488 particles | 541,233 particles | 319,989 particles | 1,145,157 particles | 223,613 particles |
| 2x iterations of 2D classification | 63,180 particles | 63,315 particles | 51,371 particles | 305,685 particles | 28,662 particles |
| Template based autopicking | 467,323 particles | 687,645 particles | 826,006 particles | N/A | 273,852 particles |
| 2D classification | 11,862 particles | 71,977 particles | 63,881 particles | 113,852 particles | 66,799 particles |
| Ab Initio 3D classification | 103,105 particles | 71,977 particles | 46,349 particles | 113,852 particles | 50,232 particles |

| **Post-merge processing** | **Combined Datasets** |
|---|---|
| 2x Iterations *ab initio* 3D classification | 405,515 particles |
| Non-uniform Refinement | 167,294 particles |
| Per-particle CTF refinement | 6.71Å map |
| Non-uniform refinement with local resolution estimation and filtering | 6.4Å map |

| Table S3: EM refinement statistics | |
|---|---|
| Resolution for refinement (Å) | 6.4 |
| $CC_{Mask}$ | 0.73 |
| RMS deviations | |
| Bond length (Å) | 0.004 |
| Bond angle (º) | 1.076 |
| No. of atoms | 8925 |
| Protein residues | 1808 |
| Ramachandran Outliers (%) | 0.4 |
| Mean B-factors ($Å^2$) | 191.4 |
| MolProbity score | 2.68 |
| Clash score | 25.11 |
